# Development of a novel murine model of in-stent neoatherosclerosis

**DOI:** 10.1101/2025.01.17.633685

**Authors:** Jiandi Liu, Demeke Geremew, Lauren Y. Sandeman, Victoria A. Nankivell, Rouyan Chen, Lei Xiang, Emma L. Solly, Liam G. Stretton, Samantha Escarbe, Stephen J. Blake, Robert A. McLaughlin, Joanne T. M. Tan, Peter J. Psaltis, Claudine S. Bonder, Jiawen Li, Yanqing Wu, Christina A. Bursill

**Affiliations:** Vascular Research Centre, Lifelong Health Theme, South Australian Health and Medical Research Institute (SAHMRI), Adelaide, South Australia 5000, Australia; Faculty of Health and Medical Sciences, University of Adelaide, Adelaide, South Australia 5005, Australia; Department of Cardiology, Second Affiliated Hospital, Jiangxi Medical College, Nanchang University, Nanchang, Jiangxi 330006, China; Australian Research Council (ARC) Centre of Excellence for Nanoscale BioPhotonics (CNBP), University of Adelaide, Adelaide, South Australia 5005, Australia; School of Electrical and Mechanical Engineering, University of Adelaide, Adelaide, South Australia 5005, Australia; Centre for Cancer Biology, University of South Australia and SA Pathology, Adelaide, South Australia 5000, Australia; Lynn Systems Immunology Laboratory, Precision Cancer Medicine Theme, South Australian Health and Medical Research Institute (SAHMRI), Adelaide, South Australia 5000, Australia; College of Medicine and Public Health, Flinders University, Adelaide, South Australia 5042, Australia; Department of Cardiology, Royal Adelaide Hospital, Adelaide, South Australia 5000, Australia

**Keywords:** stent, neoatherosclerosis, optical coherence tomography, near-infrared fluorescence imaging

## Abstract

**Objective:** In-stent neoatherosclerosis is a phenomenon of percutaneous coronary intervention with stenting. Whilst similar to *de novo* atherosclerosis, it develops rapidly over 1-5 years rather than over a lifetime. No preclinical small animal models exist that allow full elucidation of neoatherosclerosis biology and future treatments. The aim of this study was to establish and validate a novel murine model of in-stent neoatherosclerosis.

**Approach and Results:** Murine stainless-steel stents (2.5 × 0.7 mm) were deployed into donor descending aortas of atherosclerosis-prone apolipoprotein *(Apo)e*^-/-^ mice, then carotid-interposition grafted into *Apoe*^-/-^ recipients. Mice (n=6-8/group) received chow or a high cholesterol diet (HCD) for 7- or 28-days post-surgery. Multimodal intravascular imaging, simultaneously combining optical coherence tomography (OCT, plaque burden) and fluorescence for indocyanine green (ICG, plaque instability), visualized in-stent neoatherosclerosis across the entire length of the stented site. Histological analyses revealed that stented vessels from mice fed HCD had neointimas with prominent lipid cores and an elevated CD68^+^ macrophage content, similar to human neoatherosclerosis. Mice fed chow post-stenting had distinctly different neointimas that were smooth muscle cell rich, resembling neointimal hyperplasia. Consistent with this, flow cytometry revealed a higher content of monocytes/macrophages and dendritic cells in stented aortas from mice fed HCD than in non-stented aortas.

**Conclusion:** We have developed and validated the first murine model that replicates the unique characteristics of human in-stent neoatherosclerosis. This project has implications for exploring the mechanisms that promote neoatherosclerosis and testing targeted new therapies.

**RESEARCH PERSPECTIVE:** *What Is New?:*

- We have developed and validated a novel murine model of in-stent neoatherosclerosis, presenting a new platform that will facilitate the discovery of novel mechanistic targets of in-stent neoatherosclerosis and preventative therapies.
- This model develops lesions with a similar morphology to human in-stent neoatherosclerosis and distinct to in-stent neointimal hyperplasia, with higher extracellular lipid and macrophage content and proportionately less smooth muscle cells.
- We show a first-time visualization of murine in-stent neoatherosclerosis using bimodal intravascular imaging with simultaneous capture of structural information (optical coherence tomography, plaque burden) and the distribution of areas of plaque instability (high-sensitivity fluorescence, indocyanine green) within the plaque.

*What new question does this study raise?:*

- How can the utility of this novel model be maximized as a platform for discovering novel agents that prevent in-stent neoatherosclerosis?

*What question should be addressed next?:*

- Are there unique mechanisms of in-stent neoatherosclerosis, distinct to *de novo* atherosclerosis, that can be specifically targeted to prevent disease and ultimately increase stent performance?

## INTRODUCTION

Coronary artery disease (CAD), which is a narrowing or occlusion of the coronary arteries that hampers myocardial perfusion, remains the leading cause of morbidity and mortality worldwide. It is a significant cause of myocardial infarction.^1^ Coronary stenting is the primary option used by cardiologists to re-open coronary vessels blocked by atherosclerosis in an effort to restore myocardial perfusion in CAD patients.^2^ The latest-generation of drug-eluting stent (DES) has overcome the limitation of the bare metal stent (BMS) and decreased the incidence of neointimal hyperplasia-induced in-stent restenosis.^3^ However, the emergence of in-stent accelerated atherosclerosis can also burden the coronary stent. This process may occur rapidly within months to years following percutaneous coronary intervention (PCI).^4^ This pathological phenomenon is called neoatherosclerosis, which is similar to *de novo* atherosclerosis, but develops more rapidly after stent deployment rather than over a lifetime. Unlike neointimal hyperplasia that is driven by smooth muscle cell (SMC) proliferation and is SMC-rich, neoatherosclerosis is characterized by an accumulation of lipid-laden macrophage foam cells within the neointima of stented arteries, with or without necrotic core formation. Calcification or thrombosis may also occur.^4^ In-stent neoatherosclerosis has emerged as the main contributor to late in-stent restenosis and very late stent thrombosis, which are late vascular complications of stenting.^5^

The underlying mechanisms of neoatherosclerotic development are not fully elucidated, and there are no therapies that specifically target neoatherosclerosis.^5^ Thus far, the majority of preclinical studies have examined the mechanisms of neointimal hyperplasia. Consequently DES have been developed and are the most-widely used stents in clinical settings today.^6^ Very few studies have reported the development of in-stent neoatherosclerosis in animal models, however, those that do use rabbit^7,8^ and pig models.^9^ To date, no murine model exists that allows investigation of neoatherosclerotic biology and the testing of future treatments that specifically target the unique biology of neoatherosclerosis post-stenting. Murine models have the immense benefit of enabling higher throughput, lower cost, and availability of genetically modified strains, providing essential insights into the role of specific pathways in the pathophysiology of atherosclerosis and other vascular disease states.^6^ Accordingly, this study established and validated a novel murine model of in-stent neoatherosclerosis.

## METHODS

The authors declare that all supporting data are available within the article (and its Supplemental Material).

### Animals

Male atherosclerosis-prone apolipoprotein (*Apo*)*e*^-/-^ mice on a C57BL/6J background were housed in a facility with a 12-hour light/dark cycle and administered aspirin (10 mg/kg/day; Aspro Clear, Bayer, Australia) in the drinking water 1 week before surgery and throughout the study. All experimental procedures were approved by the South Australian Health and Medical Research Institute Animal Ethics Committee (SAM422.19) and conformed to the Guide for the Care and Use of Laboratory Animals (United States National Institute of Health).

### Stented Aorta-Carotid Interposition Grafting Procedure

The surgical procedure was carried out as described previously.^10^ Briefly, in the donor mouse, a stainless-steel stent (2.5 × 0.7 mm; Cambus Medical Ltd., Galway, Ireland) was crimped onto a 1.25 × 6 mm (nominal diameter × balloon length) balloon angioplasty catheter (Biotronik, Berlin, Germany). This was inserted retrograde up the descending aorta and deployed by balloon inflation to 10 atm of pressure for 30 sec. The stented vessel was harvested by electrocautery and stored in phosphate-buffered saline (PBS) at 4°C until grafting.

In the recipient mouse, the right common carotid artery was fully exposed, ligated, and divided between 7-0 silk (768G Black braided, Ethicon, NJ, USA) ties at the midpoint. Polyimide cuffs (0.6 mm diameter; Cole Parmer, IL, USA) were threaded over each end of the carotid artery and anchored by hemostatic clamps. The carotid vessels were then everted over the cuffs and secured with 8-0 silk sutures (SMOS81 Blue, Bydand Medical, Australia). The stented donor vessel was interposition-grafted into the recipient mouse by sleeving its ends over the carotid artery cuffs and securing with 8-0 silk sutures. Vessel patency was confirmed when the clamps were removed, and blood flow was restored.

Immediately following the grafting procedure, mice were fed a high cholesterol diet (HCD) containing 21% fat and 0.15% cholesterol (SF-00219, Semi-Pure Rodent Diet, Specialty Feeds, WA, Australia) or standard rodent chow for 7- or 28-days. Those mice were divided into 3 groups, including groups fed the HCD for either 1 week or 4 weeks post-surgery. A third group were fed standard chow for 4 weeks post-surgery.

At the endpoint, mice that had undergone stenting surgery were euthanized via an overdose of isoflurane anesthesia and cardiac puncture. Stented vessels and non-stented control descending aortas were excised for histological analyses or flow cytometry. Furthermore, one mouse fed HCD for 4 weeks was subjected to bimodal intravascular imaging. Plasma was isolated from whole blood by centrifugation, aliquoted and stored at -80°C until needed.

### Blood Lipids Analysis

Total cholesterol, and triglyceride levels were measured in the plasma of mice using a colorimetric assay (FUJIFILM, Osaka, Japan) according to the manufacturer’s guidelines. For high-density lipoprotein-cholesterol (HDL-C), the plasma was first incubated with 20% polyethylene glycol (1:1) for 5 mins, cholesterol was then measured on the supernatant. Low-density lipoprotein-cholesterol (LDL-C) levels were calculated as the difference between total cholesterol and HDL-C concentrations.

### Optical Coherence Tomography and Fluorescence Imaging in stented mice

Li *et al.*^11^ have designed a novel lens-in-lens intravascular imaging catheter that simultaneously acquires optical coherence tomography (OCT) and near infrared fluorescence (NIRF) images. Imaging of a stented vessel was performed in a mouse fed HCD for 4 weeks postoperatively using the novel miniaturized imaging catheter. Indocyanine green (ICG) (Diagnostic Green Ltd., Athlone, Ireland) was intravenously administered as a fluorescent agent and plaque ‘instability’ marker^12^ via the tail vein 30 minutes before the imaging procedure to allow vessel uptake. Whilst the mouse was under anesthesia, the stented vessel in the carotid region was exposed and a small incision was made at the proximal end of the vessel through which the miniaturized imaging catheter device (diameter: 0.635 mm) was inserted. Simultaneous OCT and fluorescence images were acquired as reported previously with this system.^11^ The imaging catheter was rotated and pulled back using a custom-designed automated stage at 20 µm steps until it reached the proximal end of the artery where the stent was no longer visible by OCT. After imaging, the vessel was excised for histological analysis. Three-dimensional volume rendering of the stented vessel was performed using Amira software (Thermo Fisher Scientific, USA) combining OCT and NIRF datasets. A cutaway volume-rendered view was generated to show the stented vessel lumen with neointimas, visualized using a lookup table colormap.

### JB-4 Resin Embedding of Stented Vessels

Stented vessels harvested at the end of the experimental periods were incubated with 4% phosphate-buffered paraformaldehyde (PFA) overnight at 4°C, and subsequently embedded into JB-4 (glycol methacrylate) resin (ProSciTech, Thuringowa Central, Australia) according to the manufacturer’s instructions for histomorphometry and immunohistochemistry analysis. Transverse sections (5 μm) of resin-embedded stented vessels were cut using tungsten-carbide blades on an automatic microtome (HistoCore AUTOCUT, Leica Microsystems Pty Ltd., VIC, Australia).

### Paraffin Embedding of Descending Aortas

The descending aortas of the stented mice were incubated overnight with 4% PFA at 4°C then divided evenly into three equal parts before embedding all three in the same orientation in paraffin. They were sectioned across all three parts of the aorta simultaneously at 5 μm using a semi-automated microtome (HM340E, Thermo Fisher Scientific, MA, USA).

### Histomorphometry

Histological sections were imaged using the NanoZoomer slide scanner (Hamamatsu Photonics, Hamamatsu, Japan). Immunofluorescence-stained sections were imaged using the Zeiss AxioScan Z1 slide scanner (Carl Zeiss AG, Baden-Württemberg, Germany). Image analysis of tissue sections was performed using Image Pro-Premier 9.2 software (Media Cybernetics, MD, USA). For quantification of the neointima and neointimal extracellular lipids, resin-embedded sections were stained with Multiple stain (Polysciences, PA, USA), and paraffin-embedded sections were stained with hematoxylin and eosin (H&E). Five sections spanning the length of the stented region of the vessel (or the descending aorta) were analyzed. The neointimal area per mouse was represented by calculating the average of the 5 analyzed sections. The %stenosis was equal to the average neointimal area divided by the average area inside the internal elastic lamina. The extracellular lipid content was calculated for each section and was identified as the acellular white regions in the neointima and expressed as a percentage of total neointimal area.

For quantification of the neointimal collagen and medial extracellular lipids, resin-embedded sections were stained with Masson’s Trichrome stain (ab150686, Abcam, Cambridge, UK), using the manufacturers instructions with the exception that the Biebrich scarlet-acid fuchsin solution incubation time was extended to 5 minutes and Aniline blue solution incubation time was extended to 40 minutes, as optimized for the JB-4 resin. 3-5 sections spanning the length of the stented region of the vessel were analyzed. The surface-adherent leukocytes were measured by manual cell counting. The neointimal collagen was expressed as a percentage of total neointimal area.

### Immunohistochemistry

Resin-embedded sections were subjected to antigen retrieval using a Tris-EDTA buffer (pH 9.0) and incubated with Blocking buffer (10% Goat-serum + 0.1% Tween-20 in PBS) for 6 hours. Sections were then stained for SMC α-actin using a monoclonal anti-mouse antibody conjugated to alkaline phosphatase (1:200, clone 1A4, A5691, Sigma-Aldrich Inc., MO, USA) overnight at 4°C. The Vector Red substrate kit (Vector Laboratories, Burlingame, CA, USA) was then incubated with the sections for 10 mins. Five sections spanning the length of the stented vessel (or the control descending aortas) were analyzed. The neointimal SMC content was expressed as % total neointimal area, and the medial SMC content was expressed as % total medial area.

### Immunofluorescence

Resin-embedded sections were permeabilized with Proteinase K (S3020, Dako, Agilent Technologies Inc., Santa Clara, CA, USA) and incubated with Blocking buffer (10% Goat-serum + 0.1% Tween-20 in PBS) for 6 hours. Sections were then exposed overnight at 4°C to rabbit Polyclonal anti-CD68 (1:50, 0.5 mg/mL, ab125212, Abcam, Cambridge, UK) or rabbit IgG Isotype Control (1:1100, 11.0 mg/mL, #31235, Thermo Fisher Scientific, MA, USA). Alexa Fluor^TM^ 488 Donkey anti-Rabbit IgG (H+L) Highly Cross-Adsorbed Secondary Antibody (1:500, A21206, Thermo Fisher Scientific, MA, USA) was incubated with the sections at 37°C for 2 hours, then slides were mounted in VECTASHIELD^®^ Antifade Mounting Media with DAPI (H-1200, Vector Laboratories Inc., CA, USA). Three sections spanning the length of the stented vessel were analyzed. The CD68^+^ macrophages in the neointima and vessel media (analyzed separately) was expressed as % total neointimal area or % total medial area, respectively.

### Flow Cytometry

Harvested stented vessels and non-stented descending aortas were placed into Iscove’s Modified Dulbecco’s Media (IMDM, Sigma) and digested into single-cell preparations in Hank’s Balanced Salt Solution (HBSS, Sigma-Aldrich) containing Liberase (1:100; Roche), vortexed and incubated for 45 min at 37°C. The digested homogenate was filtered through a cell strainer (70 µm). Blood samples were subjected red blood cell lysis with ammonium chloride.^13^

Aortic digests and blood samples were stained for 45 min at 4°C with fluorochrome-conjugated antibodies (Supplemental Table S1). Samples were subsequently washed with IMDM + 10% FBS and fixed with Cytofix fixation buffer (BD Biosciences). In addition, fluorescence minus one (FMO) background control samples were prepared using aortic cells from mice. Samples were run on a Cytek^®^ Aurora Spectral Analyser (Cytek^®^ Biosciences Inc.) and data analysed with FlowJo™ v10.8.1 software. Gating strategies for aortic cells and circulating cells are shown in Figures S2 and S3, respectively.

### Statistical Analysis

Data are presented as mean ± SD or as a percentage where indicated. Each data point on bar graphs represents an individual mouse. Two groups were compared using Student two-sided t-tests for parametric data or the Mann-Whitney U test for non-parametric data. When comparing more than two groups, parametric data were analyzed using one-way analysis of variance (ANOVA) with Bonferroni’s multiple comparisons or Brown-Forsythe and Welch ANOVA with Dunnett’s T3 multiple comparisons (unequal variance). Non-parametric data were analyzed using Kruskal-Wallis test with Dunn’s multiple comparisons. A *P* value of < 0.05 was considered the criterion of significance. Statistical analysis was performed using GraphPad Prism 10 software (San Diego, CA, USA).

## RESULTS

Stent deployment, using carotid grafting surgery, was conducted in 45 *Apoe*^-/-^ mice (Figure 1A), of which 36 stents were patent (no thrombosis) at the endpoint, equating to a stenting surgical success rate of 80%. The mice with in-stent thrombosis were removed from further analyses.

**Figure 1.**
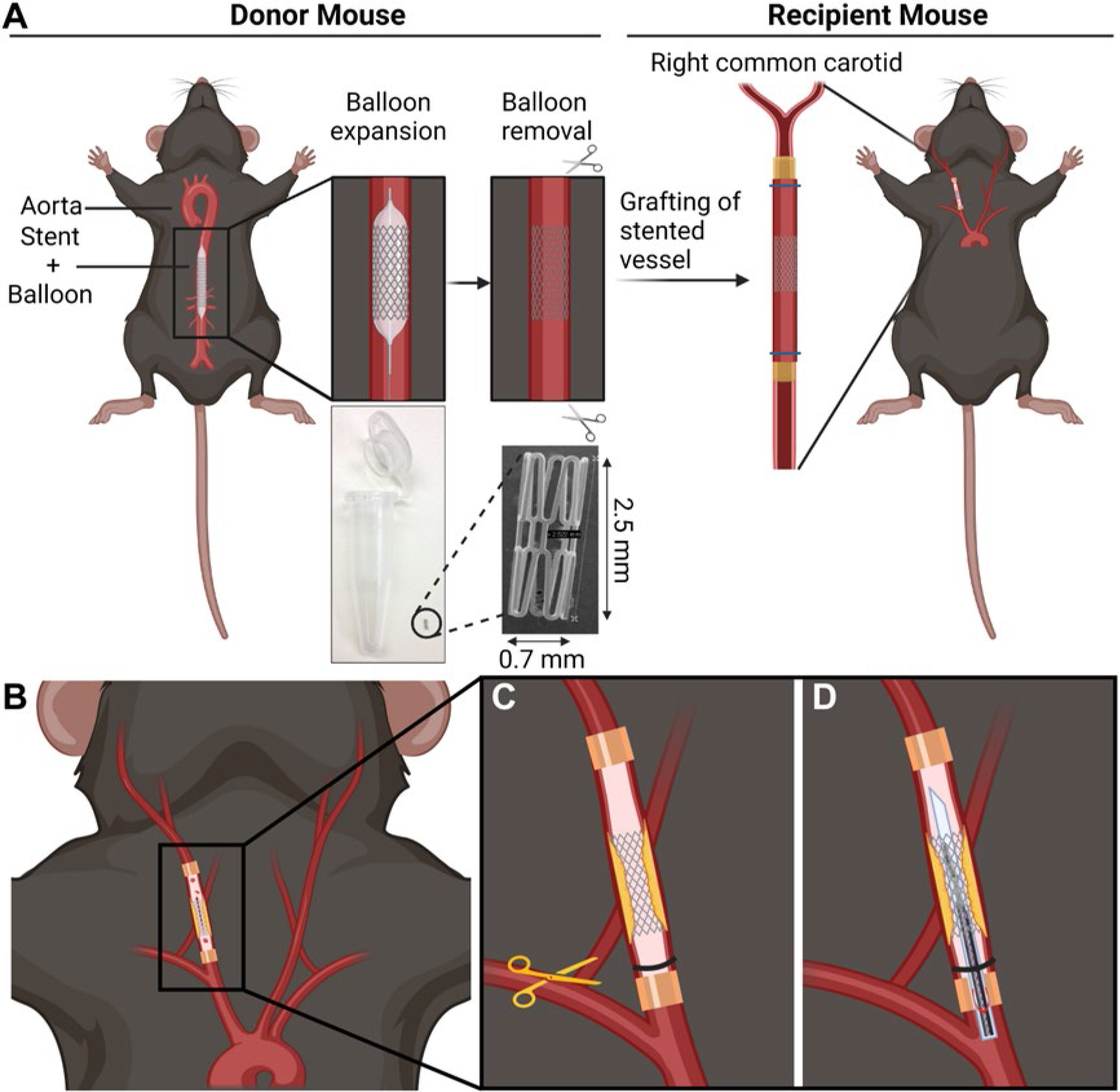
Stented Aorta-Carotid Interposition Grafting Procedure. **A**. Stainless-steel stent (2.5 × 0.7 mm) is deployed into the thoracic aorta of a donor mouse, then excised and interposition grafted into the carotid region of a recipient mouse. **B**. Under full isoflurane anesthesia, the stented vessel is fully exposed. **C**. A small incision is made at the proximal end. **D**. The imaging catheter is inserted.

### High cholesterol diet induces hypercholesterolemia in *Apoe^-/-^* mice

As expected, mice fed HCD for 4 weeks had higher average body weight, when compared to mice fed chow for 4 weeks post-surgery or HCD for 1 week, in which they did not gain weight (Table 1). Total cholesterol and LDL-C levels in mice fed HCD for 1 week and four weeks were significantly higher than in mice fed chow for 4 weeks (*P* < 0.01). Mice fed chow for 4 weeks post-surgery had higher HDL-C levels than those fed HCD for 1 week (*P*<0.01). Triglyceride levels were higher in mice fed HCD for 4 weeks than 1 week (*P* < 0.05).

**TABLE 1.** Body weight gain and plasma lipids of *Apoe*^-/-^ mice receiving stenting.

|  | 1-week HCD | 4 weeks HCD | 4 weeks Chow |
| --- | --- | --- | --- |
| No. of mice | 13 | 13 | 10 |
| Weight gain (g) | -1.06 ± 1.82 | 4.56 ± 2.55<sup>*‡</sup> | -0.18 ± 2.10 |
| TC (mmol/L) | 20.87 ± 4.02<sup>†</sup> | 28.57 ± 4.37<sup>*‡</sup> | 11.91 ± 2.80 |
| LDL-C (mmol/L) | 19.53 ± 3.67<sup>†</sup> | 26.83 ± 4.17<sup>*‡</sup> | 9.39 ± 2.55 |
| HDL-C (mmol/L) | 1.34 ± 0.70<sup>†</sup> | 1.73 ± 0.78 | 2.51 ± 0.66 |
| TG (mmol/L) | 2.65 ± 0.73 | 3.97 ± 1.18<sup>*</sup> | 3.18 ± 1.53 |

<sup>†</sup>*P* < 0.01 compared with 4 weeks Chow, <sup>\*</sup>*P* < 0.05 compared with 1 weeks HCD, <sup>‡</sup>*P* < 0.001 compared with 4 weeks Chow (Bonferroni Multiple Comparison Test). HCD, high cholesterol diet; TC, total cholesterol; LDL-C, low-density lipoprotein-cholesterol; HDL-C, high-density lipoprotein-cholesterol; TG, triglyceride.

### Visualization of in-stent neoatherosclerosis using bimodal high-resolution OCT and Fluorescence Imaging

OCT and NIRF imaging were performed simultaneously *in situ* in the stented vessel of a mouse fed HCD for 4 weeks. The mouse was injected with ICG 30 mins prior to imaging. The OCT data are shown in greyscale, bound by rings with color-coded fluorescence intensity signals that demonstrate the uptake of ICG into and across the vessel (Figure 2A-B). The OCT data are depth-resolved, providing an entire structural image, while the NIRF signal is a projection image with an accumulated measurement at each radial position. The signal intensity is represented by the colored circle, according to the color bar, where red represents the strongest signal and blue the lowest. In the OCT images, the morphology of the in-stent neointimas are visible (dashed blue line), including the location of stent struts (red triangles). As indicated by the pseudo-colored fluorescent intensity scale, the neointimal fluorescence signal at ∼12 - 3 o’clock of the dashed red line (from bottom of Figure 2B) is stronger than that in the same region of the dashed black line (from the bottom of Figure 2A). Furthermore, the matching histological morphologies of these two regions in co-registered sections are shown in Figure 2C and 2D, respectively. The histological sections also show large neointimas predominantly located on one side of the stented vessel in the ∼11 - 4 o’clock position. Calculation of neointimal extracellular lipid content (the white acellular region in the black frames) found a higher percentage in Figure 2D section than Figure 2C (10.02% vs. 6.02%). Moreover, in co-registered sections in which immunofluorescence was used to identify CD68^+^ macrophages (Figure 2E and 2F), we observed a higher proportion of CD68^+^ macrophages (in red frames) in the neointima of the stented aortic section in Figure 2F (14.1%, cross-section located at red dashed line in Figure 2G) than in Figure 2E (8.8%, cross-section located at black dashed line in Figure 2G). The IgG isotype control is presented in Figure S1, demonstrating CD68 antibody binding specificity.

**Figure 2.**
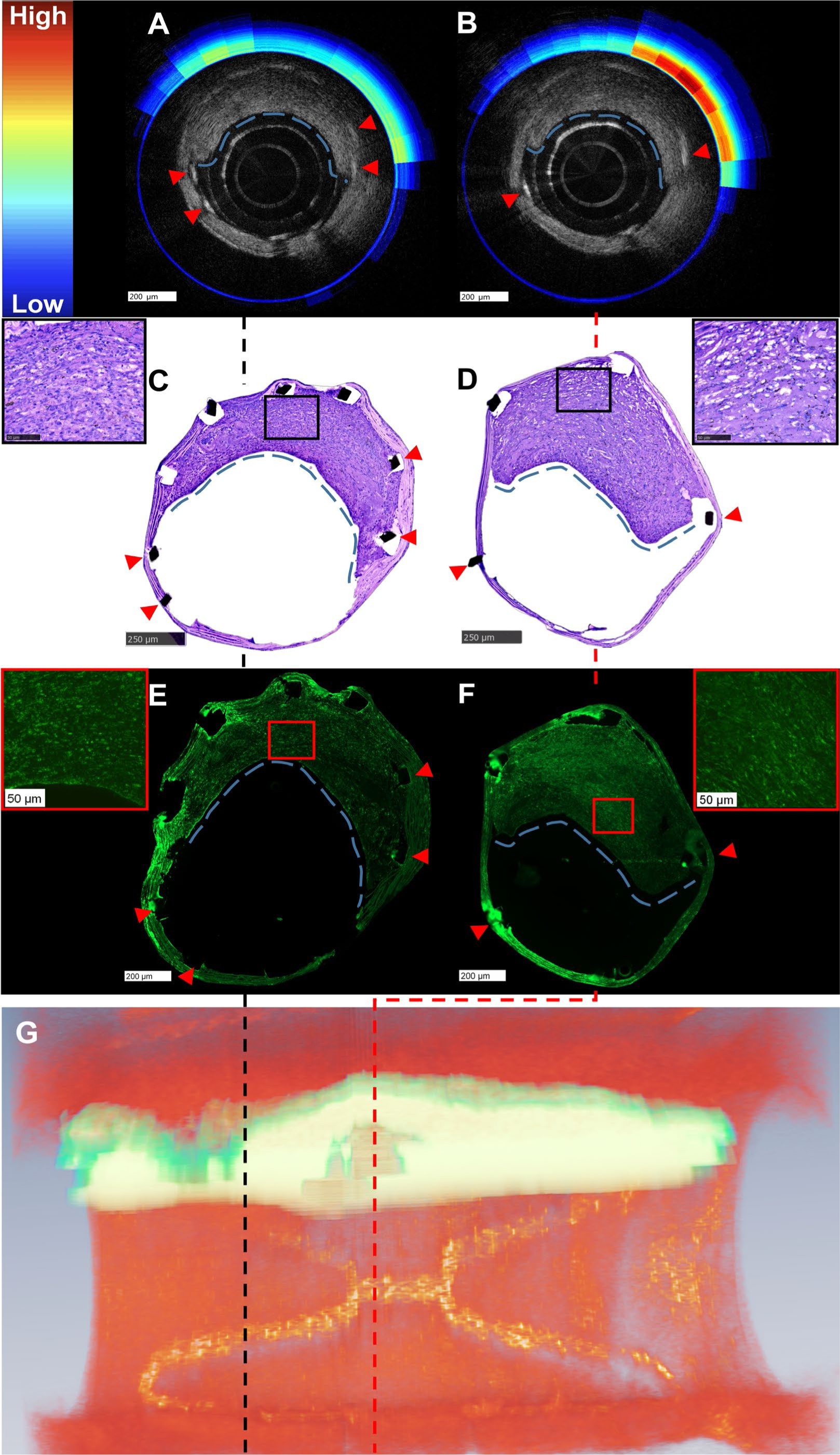
Visualization of in-stent neoatherosclerosis using bimodal high-resolution OCT and Fluorescence Imaging with 3D reconstruction. **A**. OCT (grey) and fluorescence (color) cross-sections of a stented vessel from the black dashed region of (**G**). **B**. OCT (grey) and fluorescence (color) cross-sections of the stented vessel from the red dashed region of (**G**). **C**. Co-registered histological section of (**A**, scale bars: 250 μm); zoomed images in black frame (scale bar: 50 μm). **D**. Co-registered histological section of (**B**, scale bar: 250 μm); zoomed image in black frame (scale bar: 50 μm). **E.** CD68+ immunofluorescently-stained co-registered section of (**A**, scale bar: 200 μm); zoomed image in red frame (scale bar: 50 μm). **F.** CD68+ immunofluorescently-stained co-registered section of (**B**, scale bar: 200 μm); zoomed images in red frame (scale bar: 50 μm). **G**. The 3D reconstruction of OCT+fluorescence images of the stented vessel obtained from a mouse fed HCD for 4 weeks post-stenting (also see Supplementary Video 2), red channel: OCT (plaque), light green channel: ICG fluorescence (inflammation/instability). Red triangles mark stent struts, the blue dashed line outlines neointimal edges. OCT, optical coherence tomography; HCD, high cholesterol diet; ICG, indocyanine green.

3D reconstruction of the imaging datasets allowed visualization of the plaque across the entirety of the stent segment (Figure 2G). This revealed a heterogeneous distribution of plaque along the stented vessel that included both structural features, as detected by OCT, and cellular composition by NIRF (Videos S1and S2).

### Distinct morphology of in-stent neoatherosclerosis in murine model

On resin-embedded stented aortic sections stained with Multiple stain (Figure 3A-B), we found that observationally, the stented vessels of mice fed HCD for 4 weeks, developed neointimas with foamy macrophage-like cells and extracellular lipid accumulation reminiscent of neoatherosclerosis. Even after 1 week of HCD, the neointimas of stented aortas showed evidence that lipid-laden foam cells were present (black arrowheads) and this was exaggerated further after 4 weeks of HCD. This was distinct to the morphology of mice placed on chow for 4 weeks post-surgery, in which the neointimas appear to have almost no lipid accumulation, reflecting neointimal hyperplasia. Control non-stented descending aortas displayed no neointimas as expected. Analysis of the neointimal areas revealed that there was a significant increase in neointimal area between the stented vessels from mice fed the HCD for 4 weeks, when compared to HCD for 1 week (0.10 ± 0.04 vs. 0.49 ± 0.08 mm^2^, *P* < 0.0001) (Figure 3C). There were, however, no differences in the neointimal areas of stented vessels from mice fed chow or HCD for 4 weeks. Calculation of the %stenosis revealed an increase in %stenosis between mice fed HCD for 4 weeks compared to 1 week (48.60 ± 6.44 vs. 10.89 ± 3.90 %, *P* < 0.0001) (Figure 3D). There were once again no changes in %stenosis between chow fed and 4-week HCD stented vessels. Extracellular lipid content of the neointimal areas was significantly greater within the neointimal areas in mice fed HCD for 4 weeks (18.34 ± 5.53 %) compared to 1 week of HCD (6.56 ± 2.39 %, *P* < 0.01) and 4 weeks of chow (4.75 ± 1.83 %, *P* < 0.01) (Figure 3E).

**Figure 3.**
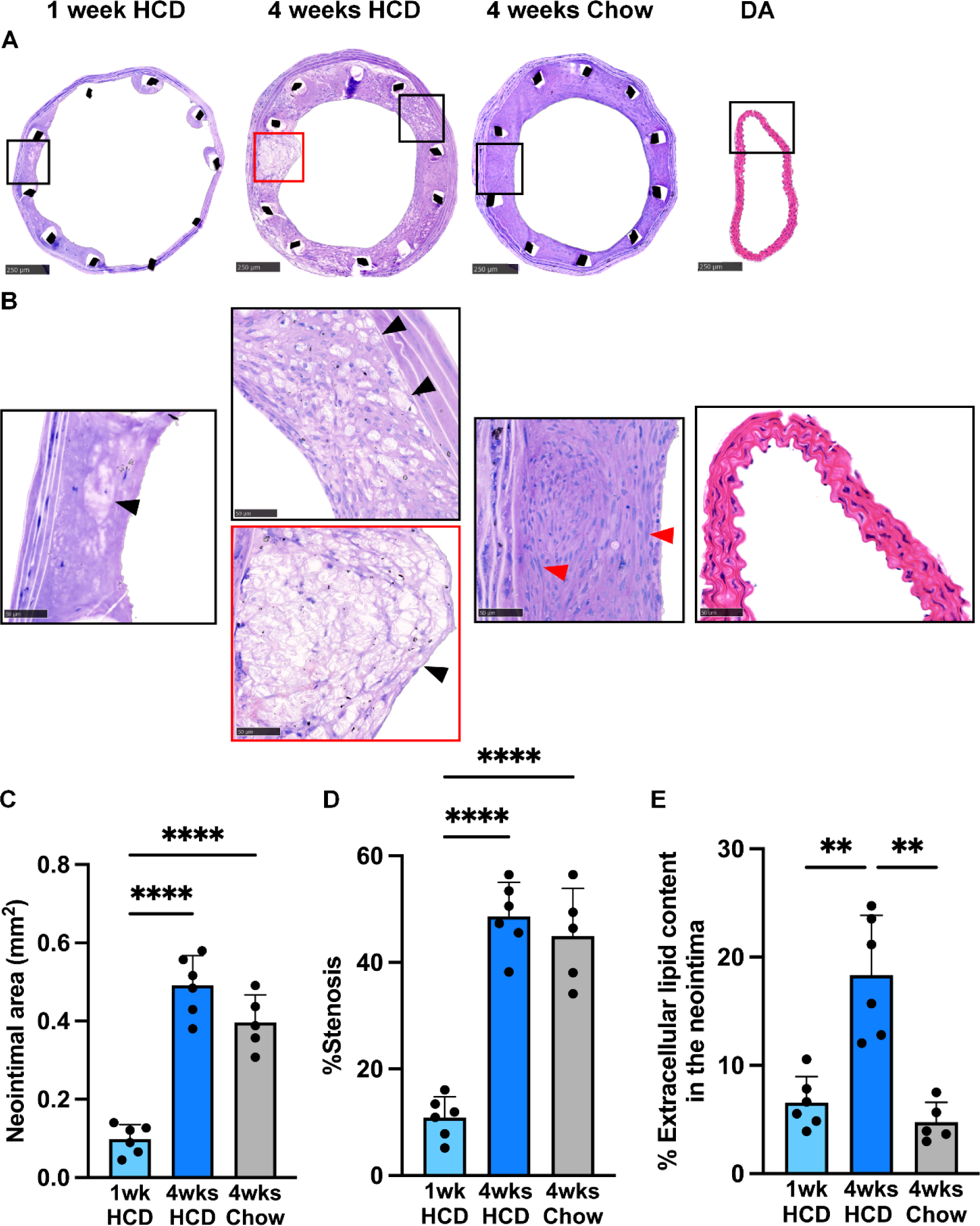
Distinct morphology of in-stent neoatherosclerosis in murine model. **A**. Representative Multiple-stained cross-sections of stented vessels grafted into *Apoe*^-/-^ mice, and haematoxylin- and eosin-stained cross-sections of non-surgical descending aortas of *Apoe*^-/-^ mice (scale bars: 250 μm). **B**. Corresponding zoomed images in the black/red frame, black triangles mark extracellular lipids (white acellular regions), red triangles mark SMCs (scale bars: 50 μm). Quantitative histomorphometry of vessel sections showing the neointimal area (**C**), percentage of stenosis (**D**), and extracellular lipid content in neointimas (**E**). Data expressed as Mean ± SD. \*\**P* < 0.01, \*\*\*\**P* < 0.0001 by one-way ANOVA with Bonferroni’s multiple comparisons or Brown-Forsythe and Welch ANOVA with Dunnett’s T3 multiple comparisons (unequal variance). HCD, high cholesterol diet; DA, descending aorta; SMCs, smooth muscle cells.

### Compositional changes between in-stent neoatherosclerosis and neointimal hyperplasia

Resin-embedded sections stained with Masson’s Trichrome demonstrated the development of distinctly different in-stent neointimas between groups (Figure 4A). Leucocytes were present near the luminal surfaces (yellow arrowheads, Figure 4B). Leukocyte number on the neointimal surface was significantly higher in mice fed HCD for 1 week (167.80 ± 124.92) compared to 4 weeks of HCD (77.33 ± 145.85, *P* < 0.05) and 4 weeks of chow (12.13 ± 4.36, *P* < 0.01) (Figure 4C). Loose collagen fibers (blue arrowheads) and lipid-laden foam cells (black arrowheads) were also present in the neointimas of stented aortas after 1 week of HCD (Figure 4D). After 4 weeks of HCD, more disorganized collagen fibers, foamy macrophage-like cells and cholesterol clefts were exaggerated in the neointimas. There was a significant increase in neointimal collagen content between 1 week and 4 weeks of HCD (27.76 ± 6.41 vs. 36.50 ± 3.65 %, *P* < 0.05) in stented vessels, yet no difference between mice fed chow and HCD for 4 weeks (Figure 4E). Interestingly, noticeable morphological changes were present in the vessel walls of stented arteries after 4 weeks of HCD, with increased presence of medial foam cells (red arrowheads, Figure 4F). Analyses of the medial lipid content revealed an increase in the mice fed HCD for 4 weeks (16.36 ± 2.52 %) compared with those fed HCD for 1 week (6.36 ± 1.57 %, *P* < 0.0001) and chow for 4 weeks (7.20 ± 3.45 %, *P* < 0.001) (Figure 4G).

**Figure 4.**
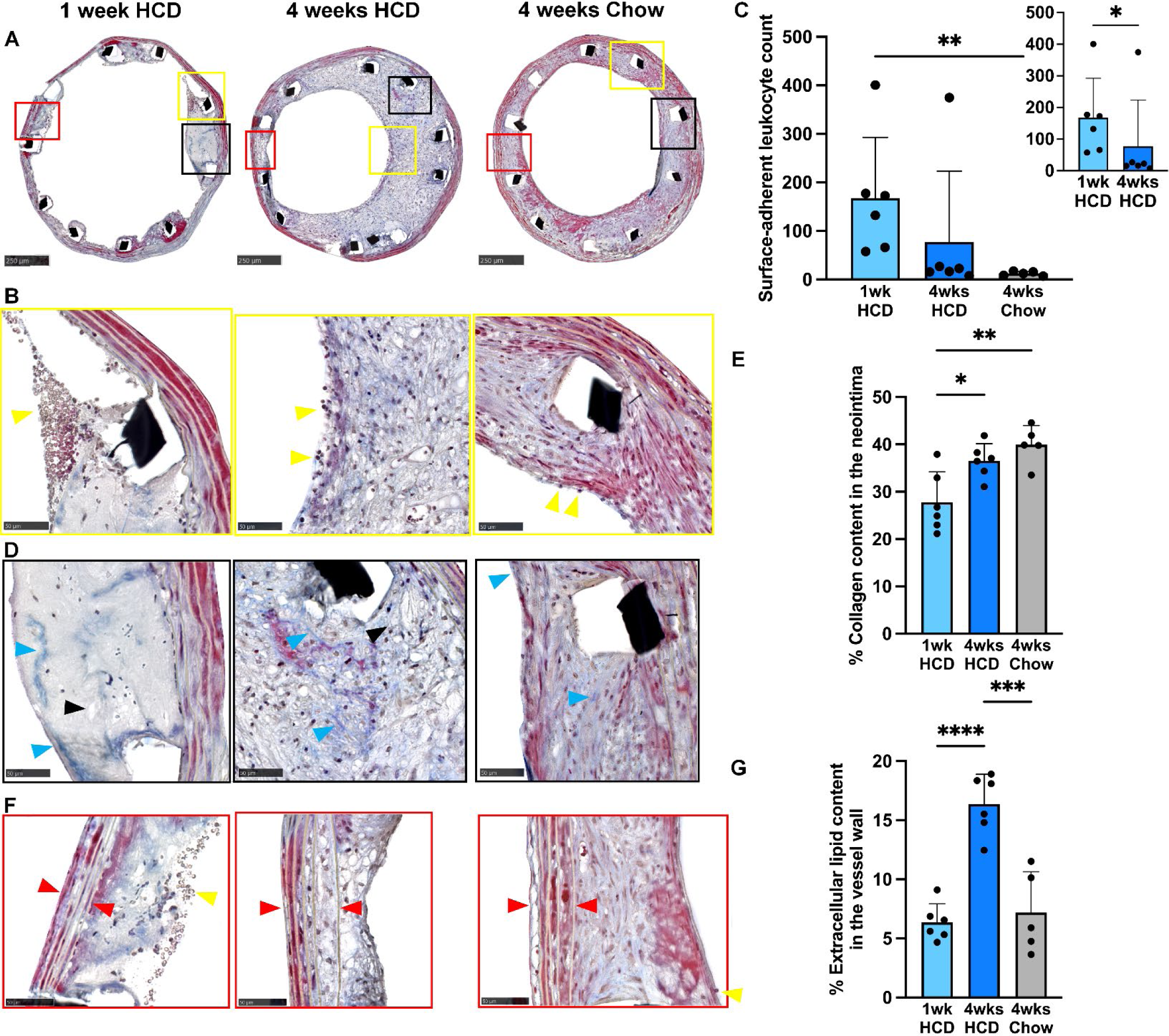
Compositional changes between in-stent neoatherosclerosis and neointimal hyperplasia. **A.** Representative Masson’s trichrome-stained cross-sections of stented vessels grafted into *Apoe*^-/-^ mice (scale bars: 250 μm). **B**. Corresponding zoomed images in the yellow frame, yellow triangles mark leukocytes (scale bars: 50 μm). **C**. Quantitative histomorphometry of vessel sections showing the surface-adherent leukocyte count. **D**. Corresponding zoomed images in the black frame, blue triangles mark collagen (blue region), black triangles mark extracellular lipids (white regions), (scale bars: 50 μm). **E**. Quantitative histomorphometry of vessel sections showing the neointimal collagen content. **F**. Higher magnification images in red frames showing medias, red triangles mark vessel walls, yellow triangles mark leukocytes (scale bars: 50 μm). **G**. Quantitative histomorphometry of vessel sections of media lipid content. Data expressed as Mean ± SD. \**P* < 0.05, \*\**P* < 0.01, \*\*\**P* < 0.001, \*\*\*\**P* < 0.0001 by one-way ANOVA with Bonferroni’s multiple comparisons (parametric data) or Kruskal-Wallis test with Dunn’s multiple comparisons (non-parametric data) or Mann-Whitney U test for comparisons between two groups (non-parametric data). HCD, high cholesterol diet.

### Reduced content of SMCs in neoatherosclerotic neointimas and medias

Immunohistochemical analysis of stented vessels revealed distinct differences in the content and distribution of α-actin^+^ SMCs in the neointimas and medias between groups (Figure 5A-C). The neointimas of stented vessels from mice fed chow for 4 weeks had the greatest content of SMCs (36.47 ± 10.91 %), and was significantly higher than that of mice fed HCD for 1 week (3.55 ± 2.96 %, *P* < 0.0001) and 4 weeks (24.31 ± 9.92 %, *P* < 0.05) (Figure 5D). In the media of the stented vessels, there were less medial SMCs in stented vessels of mice fed HCD for the 4-weeks (19.52 ± 7.92 %) than that in the 1-week HCD (45.06 ± 12.49 %, *P* < 0.01), chow-fed (39.77 ± 11.40 %, *P* < 0.05), and control non-stented descending aorta (42.18 ± 6.82 %, *P* < 0.05) groups (Figure 5E).

**Figure 5.**
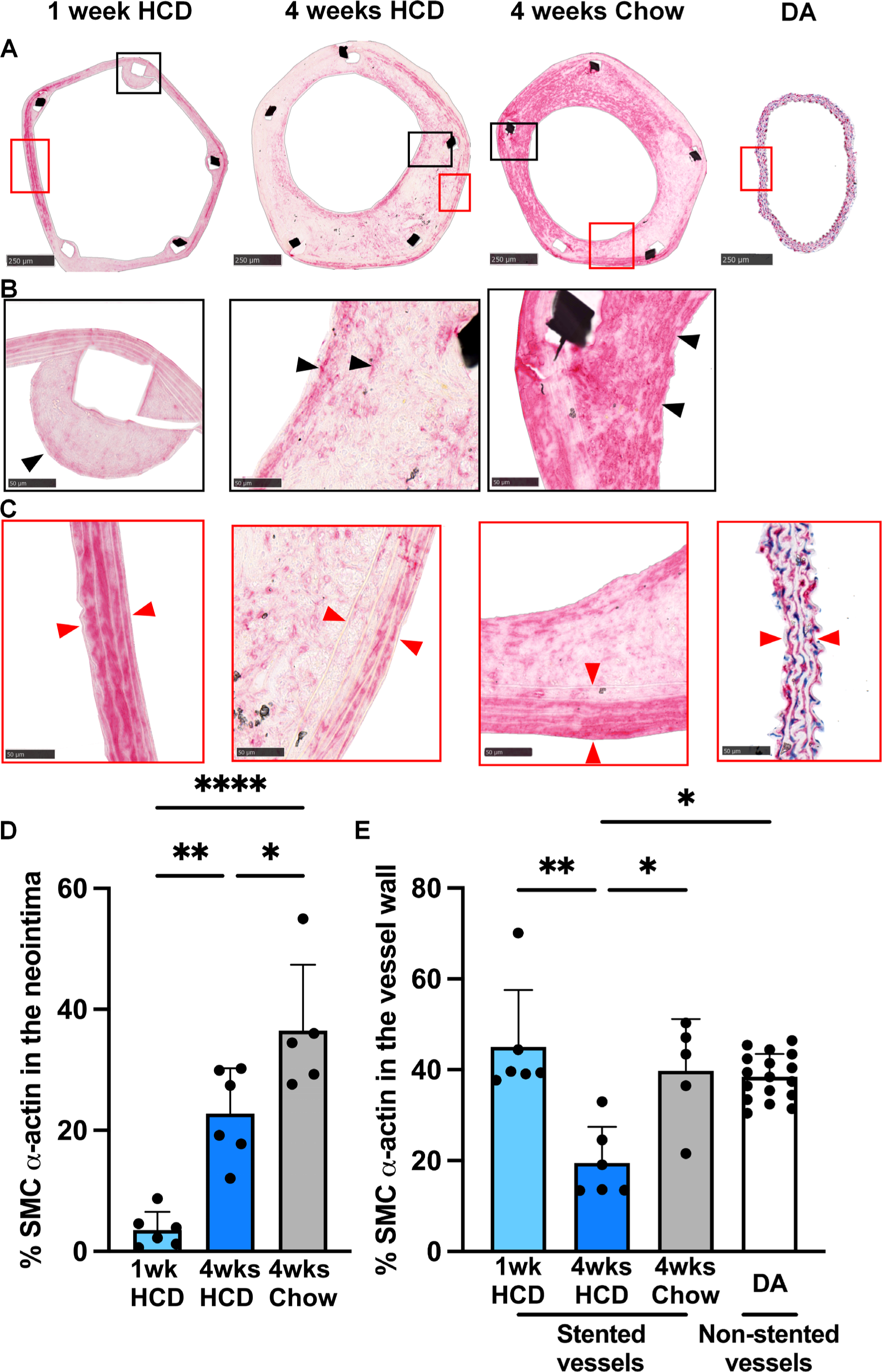
Reduced content of SMCs in neoatherosclerotic neointimas and medias. **A.** Representative α-SMA-stained cross-sections of stented vessels and non-surgical descending aortas of *Apoe*^-/-^ mice (scale bars: 250 μm). **B**. Higher magnification images in black frames, black triangles mark α-actin^+^ SMCs (red regions) in the neointima (scale bars: 50 μm). **C**. Higher magnification images in red frames, red triangles mark the vessel wall with α-actin^+^ SMCs (scale bars: 50 μm). Quantitative histomorphometry of vessel sections showing the α-actin^+^ SMC content in the neointima (**D**) and in the media (vessel wall) (**E**), respectively. Data expressed as Mean ± SD. \**P* < 0.05, \*\**P* < 0.01, \*\*\*\**P* < 0.0001 by one-way ANOVA with Bonferroni’s multiple comparisons (parametric data) or Kruskal-Wallis test with Dunn’s multiple comparisons (non-parametric data). HCD, high cholesterol diet; DA, descending aorta; α-SMA, alpha-smooth muscle actin; SMCs, smooth muscle cells.

### Increased macrophage content in in-stent neoatherosclerotic neointimas and medias

Immunofluorescence revealed the presence of macrophage-like cells in in-stent neointimas (Figure 6A-B). Macrophage content in the neointimal areas of stented vessels from mice fed HCD for 4 weeks was significantly higher (12.31 ± 3.09 %), when compared to 1 week of HCD (7.58 ± 2.61 %, *P* < 0.05) and 4 weeks of chow (6.25 ± 2.85 %, *P* < 0.05) (Figure 6C). CD68+ cells were also identified in the medias of stented vessels (Figure 6D). The CD68+ content was also greatest in the medias of stented vessels from mice fed HCD for 4 weeks (23.59 ± 6.01 %) and was statistically higher than from mice fed HCD for 1 week (11.74 ± 4.08 %, *P* < 0.01) (Figure 6E).

**Figure 6.**
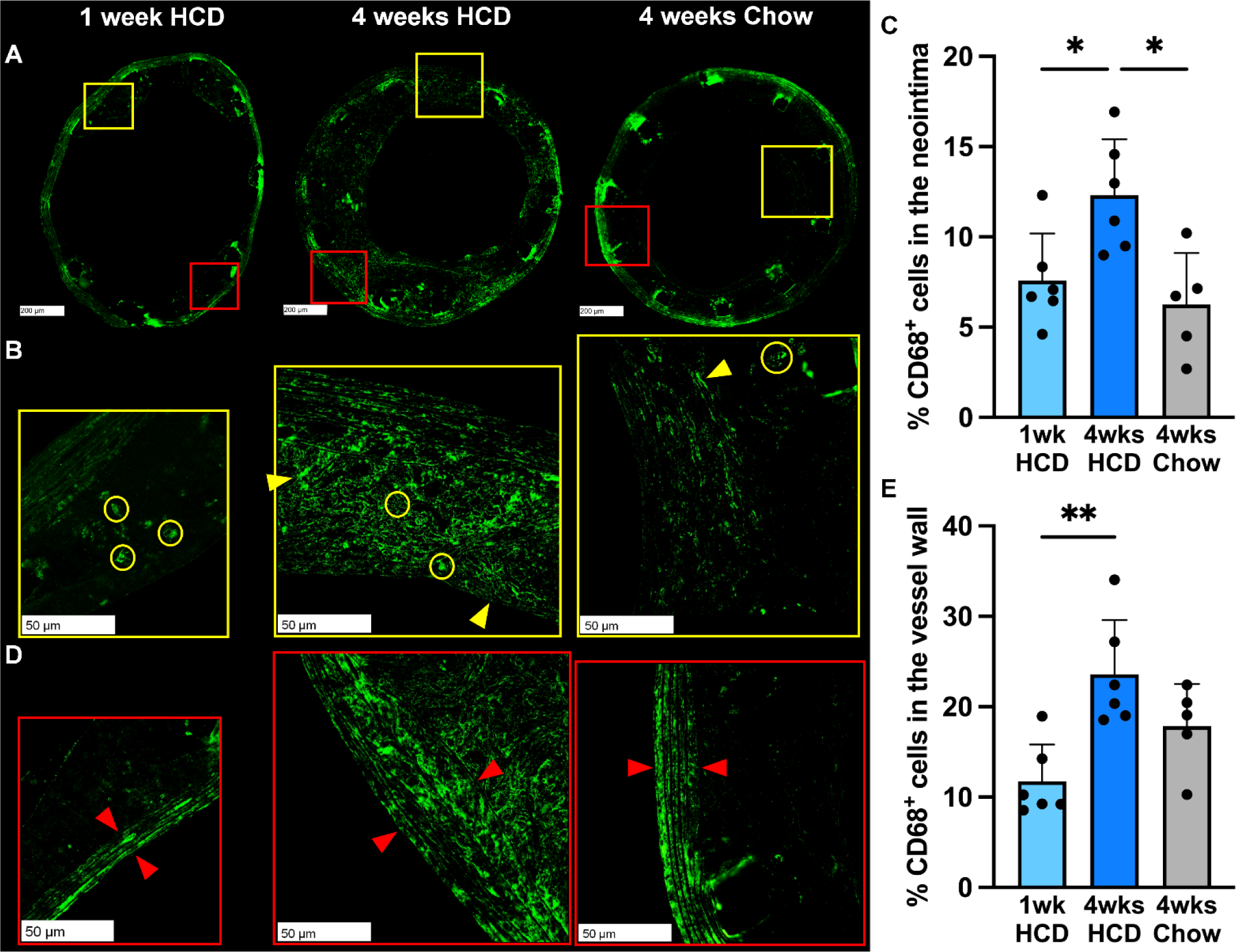
Increased macrophage content in in-stent neoatherosclerotic neointimas and medias. **A**. Representative CD68-stained cross-sections of stented vessels grafted into *Apoe*^-/-^ mice (scale bars: 200 μm). **B**. Higher magnification images in yellow frames, yellow circles and triangles mark CD68^+^ cells (green region) in the neointima (scale bars: 50 μm). **C**. Quantitative of neointimal CD68^+^ cell content. **D**. Higher magnification images in red frames, red triangles mark the vessel wall with CD68^+^ cells inside (scale bars: 50 μm). **E**. Quantitative assessment of medial CD68^+^ cell content. Data expressed as Mean ± SD. \**P* < 0.05, \*\**P* < 0.01 by one-way ANOVA with Bonferroni’s multiple comparisons. HCD, high cholesterol diet.

### Altered cellular compositions of aortas after stenting

In a separate cohort of mice, flow cytometry was utilized to characterize the cellular compositions of the stented vessels of each group. Overall, there were few statistically significant changes between groups. However, the monocyte content of stented aortas was significantly higher in mice fed HCD for 1 week (0.32 ± 0.38, *P* < 0.01), 4 weeks (0.80 ± 1.12, *P* < 0.01), and chow for 4 weeks (0.67 ± 0.66, *P* < 0.01) than non-stented descending control aortas (Figure 7A). There were no differences between groups in aortic macrophages (Figure 7B), but there were more M1 macrophages in stented aortas after 4 weeks of HCD (0.10 ± 0.08 %) and chow (0.13 ± 0.11 %) compared with non-stented aortas (0.02 ± 0.02 %, *P* < 0.05) (Figure 7C_i__-ii_). There were more M2 macrophages in stented aortas of chow-fed controls (0.08 ± 0.10 %), than that of 1 week of HCD (0.01 ± 0.01 %, *P* < 0.05) (Figure 7D). There were no changes in macrophage expression of the CD36^+^ scavenger receptor or TREM2^+^ foam cells (Figure 7E-F). There were more CD47^+^ anti-efferocytotic macrophages in stented aortas after 1 week (0.03 ± 0.02 %, *P* < 0.05) and 4 weeks of HCD (0.08 ± 0.06 %, *P* < 0.01) and 4 weeks of chow (0.07 ± 0.04 %, *P* < 0.01), than non-stented aortas (0.01 ± 0.008 %) (Figure 7G_i__-ii_). And there was a tendency toward increased CD47^+^ anti-efferocytotic macrophages in stented aortas after 4 weeks of HCD compared with that after 1 week of HCD (*P* = 0.0705) (Figure 7G_iii_). Similarly, there were more emigrating CCR7^+^ macrophages in stented aortas after 4 weeks HCD (0.02 ± 0.009 %) and 4 weeks chow (0.02 ± 0.008 %), compared non-stented aortas (0.004 ± 0.005 %, *P* < 0.05) (Figure 7H). Dendritic cells were higher in stented aortas (1-week HCD: 0.29 ± 0.27, *P* < 0.05; 4 weeks HCD: 0.60 ± 0.41, *P* < 0.05; 4 weeks chow: 0.71 ± 0.50, *P* < 0.01) than non-stented control DAs (0.12 ± 0.07 %) (Figure 7I_i__-ii_). As expected, there were significantly more endothelial cells and SMCs in control non-stented aortas, than the stented aortic groups, respectively (Figure 7J-K), but no changes in fibroblasts across all groups (Figure 7L).

**Figure 7.**
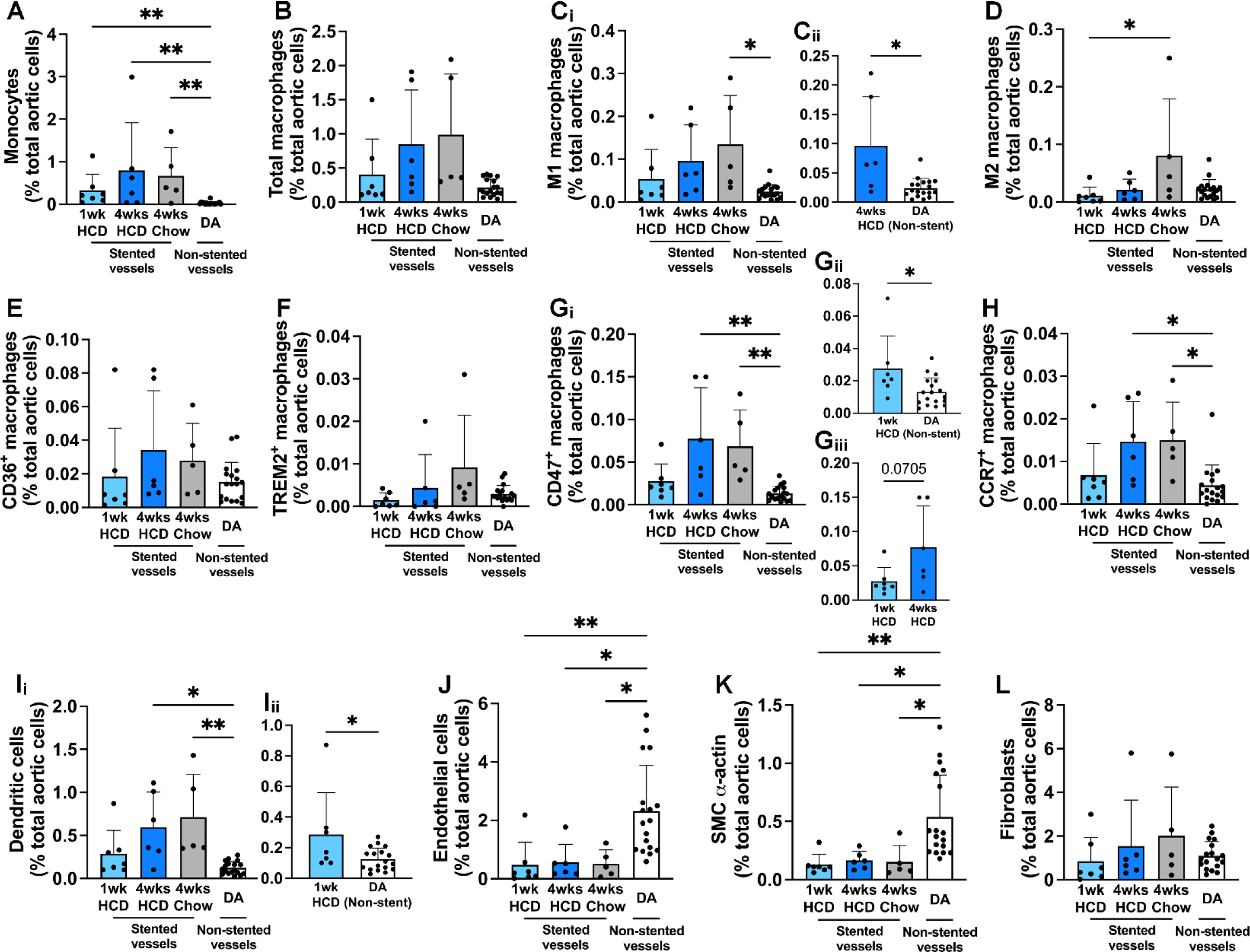
Altered cellular compositions of aortas after stenting. **A.** Monocytes, CD45^+^CD11b^+^Ly6C^+^. **B.** Macrophages, CD45^+^CD11b^+^F4/80^+^. **C.** M1 macrophages, CD45^+^CD11b^+^F4/80^+^CD86^+^CD206^-^. **D.** M2 macrophages, CD45^+^CD11b^+^F4/80^+^CD86^-^CD206^+^. **E.** CD36^+^ macrophages, CD45^+^CD11b^+^F4/80^+^CD36^+^. **F.** TREM2^+^ macrophages, CD45^+^CD11b^+^F4/80^+^TREM2^+^. **G.** CD47^+^ macrophages, CD45^+^CD11b^+^F4/80^+^CD47^+^. **H.** CCR7^+^ Macrophages, CD45^+^CD11b^+^F4/80^+^CCR7^+^. **I.** Dendritic cells, CD45^+^CD11b^+^CD11c^+^. **J.** Endothelial cells, CD45^-^CD11b^-^F4/80^-^CD31^+^. **K.** SMC α-actin, CD45^-^CD11b^-^F4/80^-^α-SMA^+^. **L.** Fibroblasts, CD45^-^CD90^+^. Data expressed as Mean ± SD. \**P* < 0.05, \*\**P* < 0.01 by Kruskal-Wallis test with Dunn’s multiple comparisons or Mann-Whitney U test for comparisons between two groups (non-parametric data). HCD, high cholesterol diet; DA, descending aorta.

### No changes in circulating immune cells

Flow cytometry on whole blood revealed there were primarily no significant differences in circulating immune cells between groups (Table 2). There were, however, higher levels of CD8^+^ T cells in mice fed chow for 4 weeks (13.44 ± 1.99, *P* < 0.05) than those fed HCD for 1 week (10.39 ± 2.32 %).

**TABLE 2.**
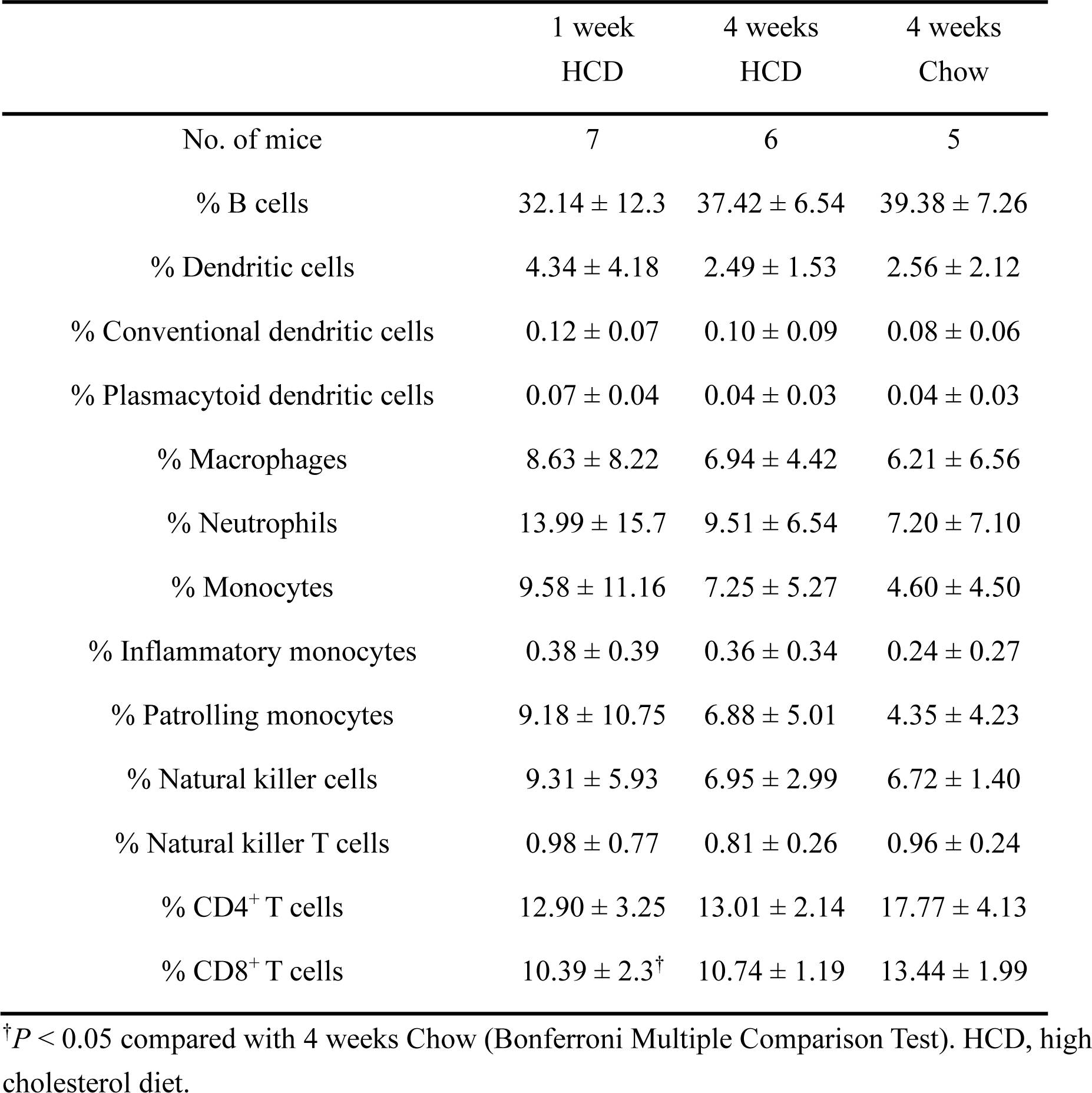
Flow cytometric quantification of circulating immune cells.

|  | 1 week<br>HCD | 4 weeks<br>HCD | 4 weeks<br>Chow |
| --- | --- | --- | --- |
| No. of mice | 7 | 6 | 5 |
| % B cells | 32.14 ± 12.3 | 37.42 ± 6.54 | 39.38 ± 7.26 |
| % Dendritic cells | 4.34 ± 4.18 | 2.49 ± 1.53 | 2.56 ± 2.12 |
| % Conventional dendritic cells | 0.12 ± 0.07 | 0.10 ± 0.09 | 0.08 ± 0.06 |
| % Plasmacytoid dendritic cells | 0.07 ± 0.04 | 0.04 ± 0.03 | 0.04 ± 0.03 |
| % Macrophages | 8.63 ± 8.22 | 6.94 ± 4.42 | 6.21 ± 6.56 |
| % Neutrophils | 13.99 ± 15.7 | 9.51 ± 6.54 | 7.20 ± 7.10 |
| % Monocytes | 9.58 ± 11.16 | 7.25 ± 5.27 | 4.60 ± 4.50 |
| % Inflammatory monocytes | 0.38 ± 0.39 | 0.36 ± 0.34 | 0.24 ± 0.27 |
| % Patrolling monocytes | 9.18 ± 10.75 | 6.88 ± 5.01 | 4.35 ± 4.23 |
| % Natural killer cells | 9.31 ± 5.93 | 6.95 ± 2.99 | 6.72 ± 1.40 |
| % Natural killer T cells | 0.98 ± 0.77 | 0.81 ± 0.26 | 0.96 ± 0.24 |
| % CD4<sup>+</sup> T cells | 12.90 ± 3.25 | 13.01 ± 2.14 | 17.77 ± 4.13 |
| % CD8<sup>+</sup> T cells | 10.39 ± 2.3<sup>†</sup> | 10.74 ± 1.19 | 13.44 ± 1.99 |

<sup>†</sup>*P* < 0.05 compared with 4 weeks Chow (Bonferroni Multiple Comparison Test). HCD, high cholesterol diet.

## DISCUSSION

The mechanisms underlying in-stent neoatherosclerosis are underexplored, which has been contributed to by the lack of validated preclinical models. The present study reports the development and validation of the first murine model of in-stent neoatherosclerosis. The surgical procedure of this model can be undertaken efficiently by a single operator, with good procedural success rates.^10^ The morphological characteristics of in-stent neoatherosclerosis in *Apoe*^-/-^ mice appear highly reproducible and are similar to other larger rabbit and porcine models as well as human neoatherosclerosis after PCI.^14^ Mice have significant advantages in that housing and breeding are cost-effective, and their size allows for smaller quantities of drug treatments or gene therapies to be used. Mechanistic assessments are possible using gene deletion, overexpression, or fluorescent labeling of specific cells or proteins that can be tracked. Furthermore, the development of atherosclerosis in *Apoe*^-/-^ mice fed HCD is relatively fast, improving modeling efficiency. Finally, there is excellent availability of antibodies, enzyme-linked immunosorbent assays, and other enzymatic or colorimetric assays for mice, which are limited for other species. Overall, our study presents a new murine model of in-stent neoatherosclerosis that will facilitate improved understanding of its mechanisms and the development and testing of novel therapeutic agents.

Our murine model development was teamed with bimodal intravascular imaging via a miniaturized imaging catheter that captured structural (i.e., OCT) and ICG fluorescence data, enabling visualization of neoatherosclerosis and the distribution of regions of instability inside the stented vessel of a mouse. Clinically available intravascular imaging techniques mostly describe gross plaque morphology without capturing critical compositional features linked to future risk.^15^ The miniaturized OCT+fluorescence dual-modality imaging catheter used in this study simultaneously acquires structural and compositional information across the entire vascular region. The fluorescence agent for dual-modality imaging was ICG, an amphiphilic near-infrared fluorophore approved by the US Food and Drug Administration as a perfusion agent.^12^ Previous studies have demonstrated that ICG can be internalized into macrophages and extracellular lipid by binding primarily to albumin and secondarily to lipoproteins. ICG is also deposited in regions of overt endothelial disruption overlying areas of macrophage infiltration and lipid-rich inflammation, rather than diffusely illuminating all areas of atherosclerosis, indicating specificity.^12,16^ Our study showed a high ICG fluorescence signal throughout the plaque, indicating the presence of lipid and macrophage-rich plaque, which was confirmed by histology. Together, this novel technology is very promising for in-stent intravascular imaging, allowing the operator to simultaneously obtain additional information about the target vessel, which is of great significance for preventing postoperative complications of PCI.

We verified this model predominantly using histology. In mice fed HCD after stenting, lipid-laden cells and a fatty streak appearance were evident in the in-stent neointimas after 1 week. This was exacerbated after 4 weeks, with a large number of foamy cells, forming typical lesions of in-stent neoatherosclerosis. By contrast, the neointimas of stented vessels from chow fed mice were distinctly different and rich in SMCs with little extracellular lipid. This was verified using histological analyses and consistent with previous studies.^10^ This highlights that the addition of HCD for relatively short periods of time post-stenting causes accumulation of lipids and drives the development of human-like in-stent neoatherosclerosis. Furthermore, the luminal surface showed a large number of endothelial-adherent leukocytes 1 week after stenting, indicative of an early-stage inflammatory response with leukocyte recruitment, which decreased after 4 weeks.^17^ Collagen content in the neointima is a determinant of plaque stability, also increased over time post-stenting, consistent with previous studies in human stented samples.^18^ SMCs also play a role in stabilizing plaques. SMCs migrate from the media to the luminal surface in response to plaque expansion to promote cap formation.^19^ Similarly, in our model, there were SMCs on the luminal surface of the in-stent neointimas in mice fed HCD for 4 weeks. This was distinct to the chow-fed group in which SMCs were prominent throughout the neointimas, demonstrating a morphology typical of neointimal hyperplasia.

We found CD68^+^ cells in the in-stent neointimas of mice fed HCD for 4 weeks that were spherical and spindle-shaped, representing SMC-derived macrophages. SMC-derived macrophages (CD68^+^) serve as the leading source of foamy macrophages in atherosclerosis,^20,21^ which lose their contractile function and acquire a phagocytic function similar to macrophages.^22^ Furthermore, the media of stented vessels after 4 weeks of HCD also featured many foam-like cells with an overall lower α-actin^+^ expression and, conversely, a higher CD68^+^ expression. These findings are consistent with previous studies that show cholesterol loading SMCs rapidly promotes transdifferentiation to a macrophage-like foam cell with elevated expression of macrophage-related genes, while losing the expression of SMC markers, including α-actin and myosin heavy chain.^23^ In the present study, we suggest that 4 weeks of HCD promoted more cholesterol into the subendothelial area, subsequently inducing more medial SMCs to transdifferentiate into macrophage-like foam cells.^24^ Taken together, our findings provide further evidence that our novel murine model is representative of a human-like neoatherosclerosis.

Flow cytometry was performed to further investigate the cellular composition of stented aortas. Overall, there were few differences between the three stented vessels groups. However, we did find striking differences in cell compositions between the non-stented aortas and the stented aortas, as expected. There were more monocytes, M1-like macrophages, efferocytotic CD47^+^ macrophage and emigrating CCR7^+^ macrophages, as well as dendritic cells in stented vessels than non-stented vessels. Stent placement causes local inflammation and blood flow disturbance associated with the overexpression of adhesion molecules. This promotes infiltration of circulating monocytes into the subendothelial space, where they convert into macrophages.^14^ We found a higher number of M1-like macrophage in stented aortas compared to non-stented aortas. M1-like macrophages are key contributors to in-stent stenosis and not only promote foam cell formation and the development of fibroatheroma with necrotic cores,^24^ but also stimulate SMC proliferation via miR-222 originating from M1 macrophage-derived exosomes.^25^ The upregulation of CD47 expression on macrophages interrupts efferocytosis and can cause accumulation of more macrophages in the neointimas of stented aortas.^26^ When macrophage content increases, protective mechanisms to reduce macrophage number are triggered, including increased macrophage egress via upregulation of CCR7 and its ligands CCL19 and CCL21 into the local draining lymph nodes.^27^ The increase in macrophage CCR7 expression found in our study in stented vessels suggests a feedback response to the expanding neointimas. Dendritic cell accumulation can not only trigger T cell activation, proliferation, and the release of pro-atherogenic cytokines (i.e., IFN-γ and TNF-α) but also accumulate intracellular lipids and transform into CD11c^+^ foam cells.^28^ Dendritic cells and vascular SMCs can interact with reciprocal stimulation, possibly perpetuating inflammation and neointimal hyperplasia.^29^ Furthermore, our flow cytometry data provides evidence that stenting damages the endothelium, as there were significantly less endothelial cells in stented vessels than the control non-stented vessels, as expected.^30,31^ We did not find differences in circulating immune cells, suggesting stenting and HCD have little effect on them and that the development of in-stent neointimas depends on local macrophage and SMCs proliferation rather than the recruitment of peripheral immune cells.^32,33^

Our study has some limitations. The surgery requires a high level of microsurgical skill and despite aspirin administration, stent thrombosis occurred in 17.8% of stented vessels. The use of additional antiplatelet agents, such as clopidogrel, may further decrease the thrombosis rate, but dual antiplatelet therapy carries the risk of increased bleeding post-surgery.^34^ Furthermore, we induced neoatherosclerosis by feeding mice a HCD after stenting and did not stent directly onto existing plaque. We believe our approach was cleaner in developing this model for the first time, however, future studies stenting on preexisting plaque will further simulate what happens clinically and the pathology of human in-stent neoatherosclerosis.

In conclusion, we have developed the first murine model that replicates the unique characteristics of human in-stent neoatherosclerosis. This project has implications for exploring the mechanisms that promote neoatherosclerosis and testing novel therapies that specially target this emerging phenomenon of PCI.

## Nonstandard Abbreviations and Acronyms

CAD: coronary artery disease
DES: drug-eluting stent
BMS: bare metal stent
PCI: percutaneous coronary intervention
SMC: smooth muscle cell
HCD: high cholesterol diet
HDL-C: high-density lipoprotein-cholesterol
LDL-C: low-density lipoprotein-cholesterol
OCT: optical coherence tomography
NIRF: near infrared fluorescence
ICG: indocyanine green
PFA: phosphate-buffered paraformaldehyde
DCs: dendritic cells

## Acknowledgments

Not applicable

## Supplemental Material

**Table S1.** Flow cytometry antibodies. Antibodies for flow cytometry analyses of aortas and blood. α-SMA, α-smooth muscle actin; APC, allophycocyanin; FITC, fluorescein isothiocyanate; PE, phycoerythrin.

| Marker | Fluorophore | Concentration | Supplier |
| --- | --- | --- | --- |
| Aorta |  |  |  |
| L/D | Zombie NIR™ | 1:000 | BioLegend |
| Ly6C | PerCP-Cy5.5 | 1:50 | BD Biosciences |
| CD45.2 | BUV395 | 1:50 | BD Biosciences |
| CD11b | APC-Cy7 | 1:50 | BD Biosciences |
| CD36 | PECy7 | 1:50 | BD Biosciences |
| CD11c | PE-CF594 | 1:50 | BD Biosciences |
| CD31 | BV421 | 1:50 | BD Biosciences |
| F4/80 | BV785 | 1:50 | BioLegend |
| CD206 | AF647 | 1:50 | BD Biosciences |
| CD86 | BV650 | 1:50 | BD Biosciences |
| CCR7 | BUV737 | 1:50 | BD Biosciences |
| CD90 | PE | 1:50 | BD Biosciences |
| α-SMA | GFP | 1:50 | Sigma |
| CD47 | BV711 | 1:50 | BD Biosciences |
| TREM2 | AF700 | 1:50 | R&D Systems |
| Blood |  |  |  |
| Viability | Zombie NIR™ | 1:1000 | BioLegend |
| CD19 | PerCP-Cy5.5 | 1:100 | BD Biosciences |
| CD45.2 | BUV737 | 1:100 | BD Biosciences |
| CD11b (M1/70) | BUV395 | 1:100 | BD Biosciences |
| F4/80 | BV785 | 1:50 | BioLegend |
| B220 | PE-CF594 | 1:500 | BD Biosciences |
| CD11c | BV421 | 1:200 | BD Biosciences |
| CD3e | R718 | 1:100 | BD Biosciences |
| CD4 | BV510 | 1:200 | BD Biosciences |
| CD8 | BV650 | 1:200 | BD Biosciences |
| NK1.1 | APC | 1:200 | BioLegend |
| MHCII | BV711 | 1:500 | BioLegend |
| Ly6G | PeCy7 | 1:400 | BD Biosciences |
| CCR2 | PE | 1:100 | BD Biosciences |
| Ly6C | BV605 | 1:100 | BD Biosciences |

---

**Figure S1.**
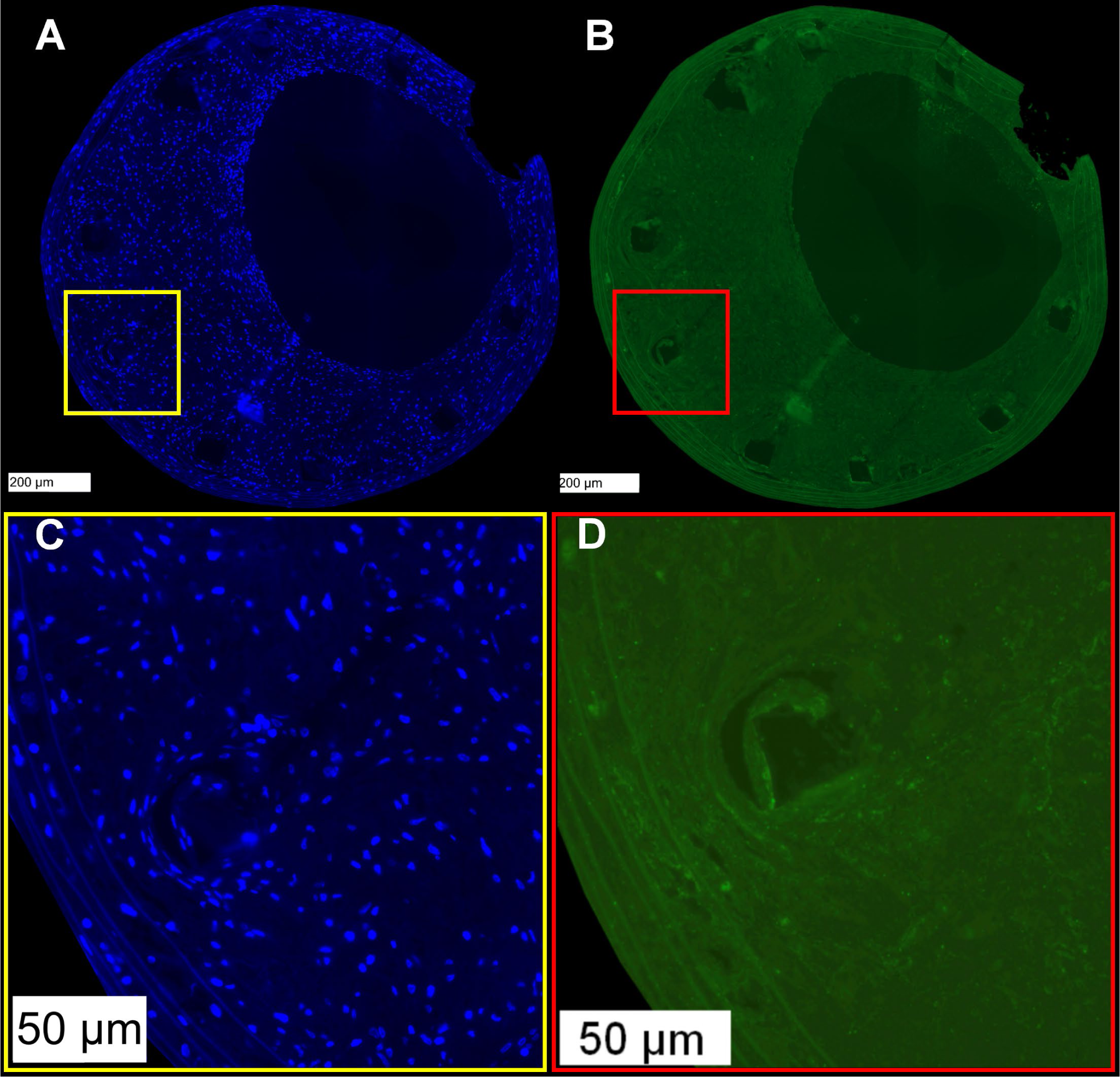
CD68 Immunofluorescence IgG control. Rabbit IgG Isotype Control (1:1100, 11.0 mg/mL, #31235). **A,** the DAPI-stained cross-section of stented vessels grafted into Apoe^-/-^ mice, scale bars: 200 μm. **B,** Rabbit IgG-stained cross-sections of stented vessels grafted into Apoe^-/-^ mice, scale bars: 200 μm. **C,** Corresponding zoomed images in the yellow frame, nuclei are visualized (blue), scale bars: 50 μm. **D,** Corresponding zoomed images in the red frame, showing the background.

**Figure S2.**
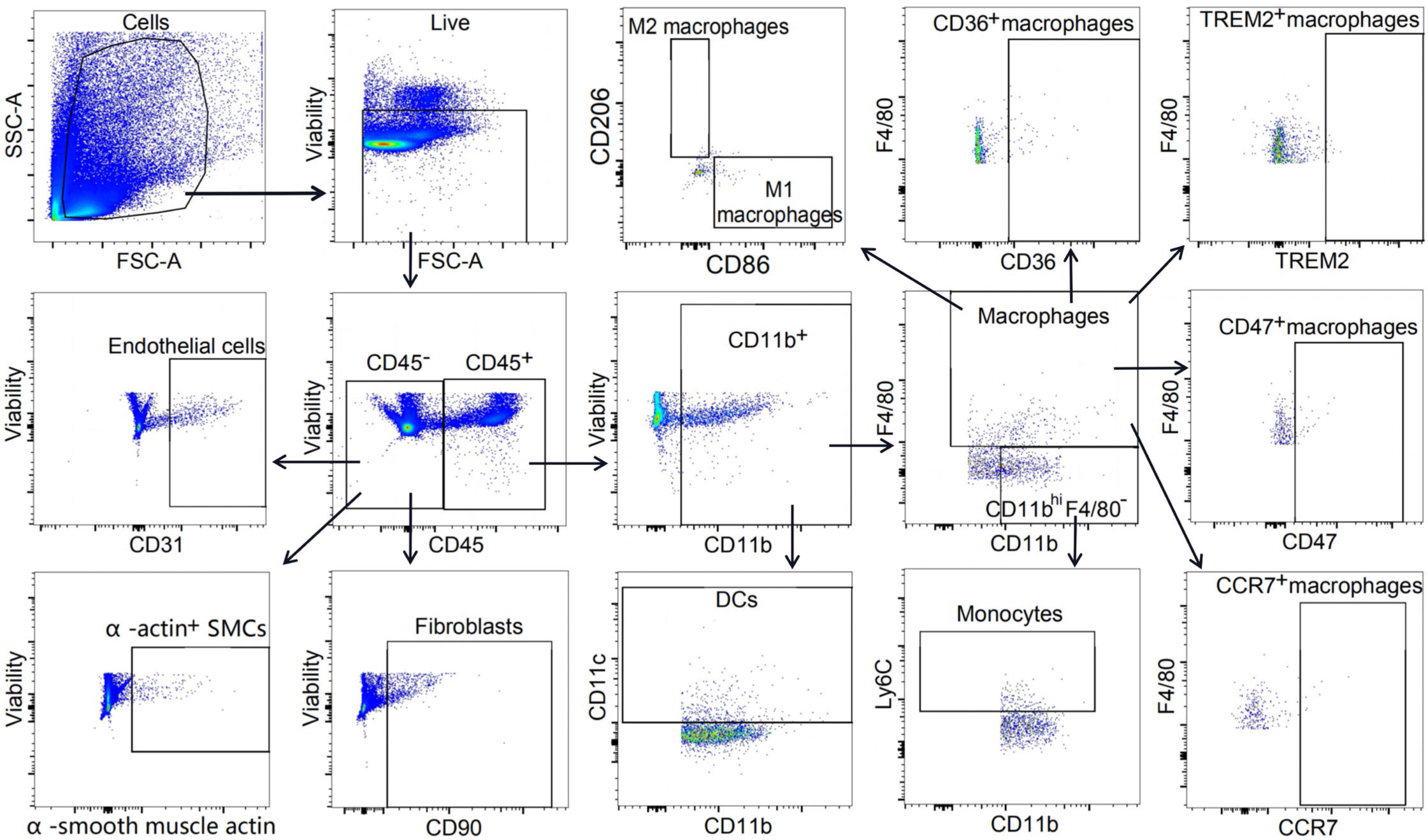
Gating strategy for murine aortic cells. Excised descending thoracic aortas and aortic arches were digested into single-cell suspensions prior to incubation with fluorescently labelled antibodies. They were then subjected to flow cytometry assessment. SMCs, α-smooth muscle cells; DCs, dendritic cells.

**Figure S3.**
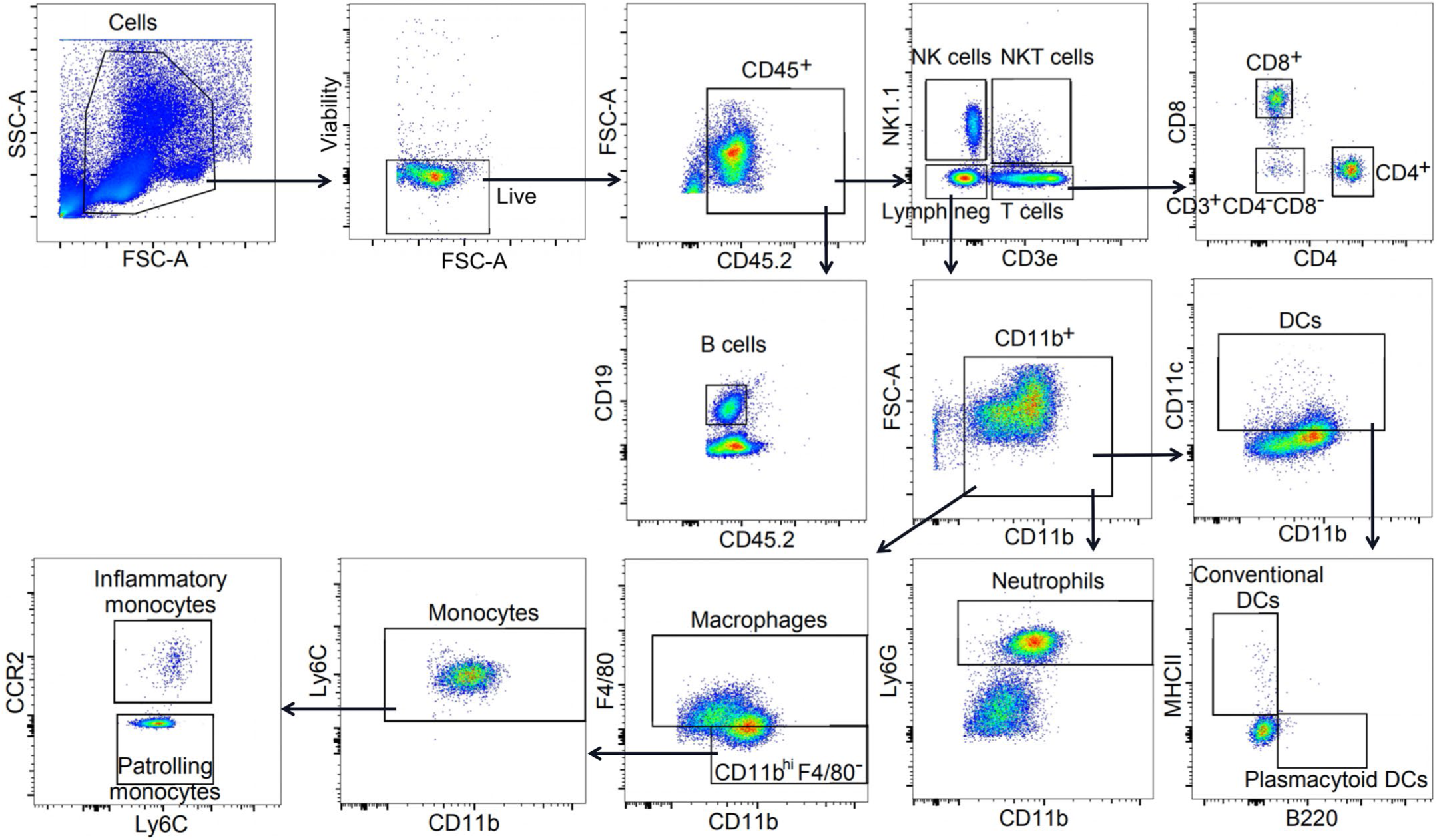
Gating strategy for circulating immune cells. Whole blood was incubated with fluorescently labelled antibodies and the subjected to flow cytometry analyses. Lymph neg, lymphocyte negative; NK, natural killer; DCs, dendritic cells.

## Figure legends for supplementary videos

**Video S1. Bimodal intravascular optical coherence tomography and near infrared fluorescence imaging of murine in-stent neoatherosclerosis.** Stents were deployed into the descending thoracic aortas of donor mice that were then excised and carotid-interposition grafted in recipient mice who were then place on high-cholesterol diet for four weeks. Indocyanine green (ICG) was injected intravenously before optical coherence tomography (OCT) and near infrared fluorescence (NIRF) imaging were performed simultaneously *in situ* across the entire stented region of the vessel. OCT images are in greyscale, bound by rings with color-coded fluorescence intensity signals that demonstrate the uptake of ICG into and across the vessel.

**Video S2. 3D visualization of bimodal optical coherence tomography and near infrared fluorescence images, taken intravascularly of murine in-stent neoatherosclerosis.** A 3D reconstruction of optical coherence tomography (OCT) and near infrared fluorescence (NIRF) data sets was generated of murine in-stent neoatherosclerosis.

